# Integrated single-cell sequencing reveals principles of epigenetic regulation of human gastrulation and germ cell development in a 3D organoid model

**DOI:** 10.1101/2022.02.10.479957

**Authors:** Alex Chialastri, Eyal Karzbrun, Aimal H. Khankhel, Monte J. Radeke, Sebastian J. Streichan, Siddharth S. Dey

**Affiliations:** Department of Chemical Engineering, University of California Santa Barbara, Santa Barbara, CA 93106, USA; Center for Bioengineering, University of California Santa Barbara, Santa Barbara, CA 93106, USA; Kavli Institute for Theoretical Physics, Santa Barbara, CA 93106, USA; Department of Physics, University of California Santa Barbara, Santa Barbara, California 93106, USA; Biomolecular Science and Engineering, University of California, Santa Barbara, CA 93106, USA; Neuroscience Research Institute, University of California Santa Barbara, Santa Barbara, 19 CA 93106, USA

## Abstract

The emergence of different cell types and the role of the epigenome in regulating transcription is a key yet understudied event during human gastrulation. Investigating these questions remain infeasible due to the lack of availability of embryos at these stages of development. Further, human gastrulation is marked by dynamic changes in cell states that are difficult to isolate at high purity, thereby making it challenging to map how epigenetic reprogramming impacts gene expression and cellular phenotypes. To overcome these limitations, we describe scMAT-seq, a high-throughput one-pot single-cell multiomics technology to simultaneously quantify DNA methylation, DNA accessibility and the transcriptome from the same cell. Applying scMAT-seq to 3D human gastruloids, we characterized the epigenetic landscape of major cell types corresponding to the germ layers and primordial germ cell-like cells (hPGCLC). As the identity of the progenitors that give rise to human PGCLCs remain unclear, we used this system to discover that the progenitors emerge from epiblast cells and show transient characteristics of both amniotic- and mesoderm-like cells, before getting specified towards hPGCLCs. Finally, as cells differentiate along different lineages during gastrulation, we surprisingly find that while changes in DNA accessibility are tightly correlated to both upregulated and downregulated genes, reorganization of gene body DNA methylation is strongly related to only genes that get downregulated, with genes that turn on displaying a lineage trajectory-dependent correlation with DNA methylation. Collectively, these results demonstrate that scMAT-seq is a high-throughput and sensitive approach to elucidate epigenetic regulation of gene expression in complex systems such as human gastrulation that are marked by rapidly transitioning cell states.

## Introduction

Regulation of gene expression in mammalian systems is tightly regulated by several layers of the epigenome that ensure precise cell type specific programs^1^. Therefore, mapping the genomewide epigenetic landscape of critical features such as DNA accessibility and DNA methylation (5-methycytosine or 5mC) is central to understanding how these regulatory factors tune complex cellular phenotypes in dynamic systems such as early post-implantation mammalian development. While the role of epigenetic reprogramming in mouse gastrulation has been extensively studied, it remains unclear how reorganization of DNA accessibility and 5mC are coupled with the emergence of cell types during human gastrulation^2^. Further, most current methods for quantifying these epigenetic features rely on the ability to isolate the desired cell type at high purity that is achieved either using cell type specific fluorescent reporters in genetically modified organisms or through access to high-quality cell type specific antibodies. However, the former approach cannot be extrapolated to humans, and many cell types also lack well-defined and unique cell surface markers and/or high-quality antibodies. Furthermore, antibodies can fail to capture transient cell states that do not present the necessary antigen, and thus these approaches are not ideally suited for studying complex systems like early human embryogenesis, that is characterized by a series of rapidly transitioning cell states^3^.

## Results

To overcome these limitations, we present a single-cell multiomics method scMAT-seq to simultaneously quantify DNA <u>m</u>ethylation, DNA <u>a</u>ccessibility, and the <u>t</u>ranscriptome from the same cell, thereby providing a marker-free approach to map the epigenetic landscape during human gastrulation using an organoid model of development. While two recent methods have been developed to make all three measurements from the same cell, these techniques rely on the physical separation of DNA and RNA prior to amplification, resulting in low throughput and a potential loss of material, limiting its usefulness^2,4,5^. To resolve this, single cells in scMAT-seq are sorted into 384-well plates and the following two steps are performed simultaneously – mRNA is reverse transcribed using a poly-T primer with an overhang containing a cell- and mRNA-specific barcode, a unique molecule identifier (UMI), the 5’ Illumina adapter and a T7 promoter; and the methyltransferase M.CviPI is used to methylate cytosines in a GpC context within open chromatin (Fig. 1a). Performing these two steps simultaneously is critical to minimize mRNA degradation and to ensure that the *in vivo* state of chromatin can be captured immediately after cell lysis. Next, second strand synthesis is used to generate cDNA, chromatin is stripped off gDNA using proteases, and 5-hydroxymethycytosine (5hmC) sites in the genome are glucosylated to block downstream detection by the restriction enzyme MspJI. Thereafter, MspJI is added, which recognizes methylated cytosines in the genome and creates double-stranded DNA breaks that are ligated to adapters containing a cell- and gDNA-specific barcode, a UMI, the 5’ Illumina adapter and a T7 promoter^6^. Following this step, all molecules are tagged with cell- and molecule-of-origin-specific barcodes, and contain a T7 promoter, allowing samples to be pooled and amplified by *in vitro* transcription (IVT). As described previously, Illumina libraries are then prepared, enabling simultaneous quantification of mRNA, 5mC and DNA accessibility from the same cell without requiring physical separation of the nucleic acids^6–8^. Finally, depending on the context of the methylated cytosine in the sequencing data, gDNA reads are either assigned to the methylome (CpG context) or DNA accessibility (GpC context) dataset for each individual cell^2,4,5,9^.

**Figure 1.**
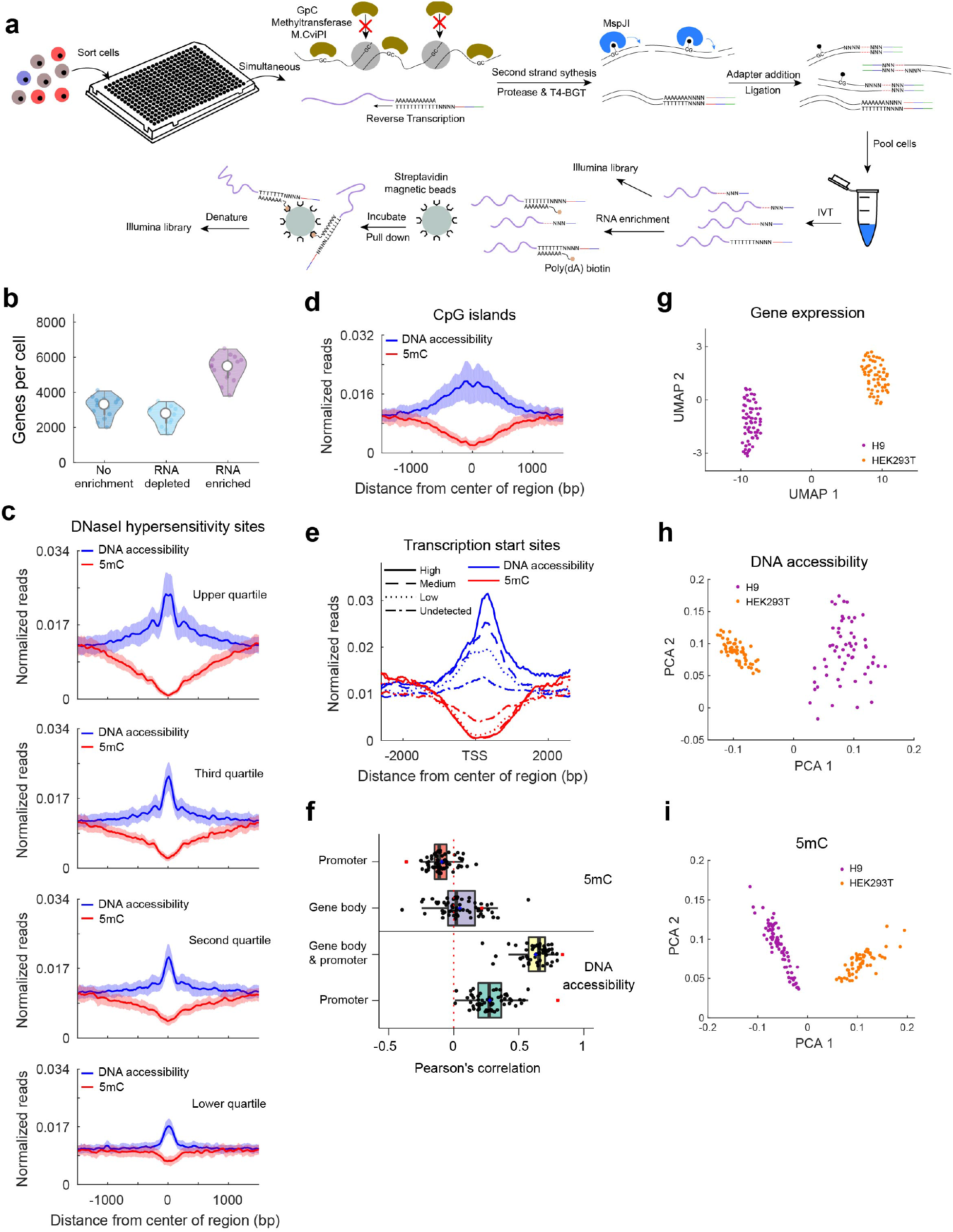
Joint profiling of DNA methylation, DNA accessibility, and the transcriptome from the same cell using scMAT-seq. (**a**) Workflow of scMAT-seq, where the first step involves simultaneous reverse transcription of mRNA and marking of open chromatin with M.CviPI. After second strand synthesis, protease and T4-BGT treatment, methylated cytosines in the genome are digested using MspJI. The barcoded cDNA and gDNA molecules are then pooled and amplified using IVT. This is followed by mRNA enrichment and Illumina library preparation. In the schematic, RNA is shown in purple, DNA in black, cell- and mRNA-/gDNA-specific barcodes in red, Illumina read 1 sequencing primer in blue, and T7 promoter in green. (**b**) Violin plot shows the number of genes detected per cell with or without RNA enrichment. Dots represent individual cells. (**c**,**d**) Averaged single-cell DNA accessibility (blue) and DNA methylation (red) profiles at DNase I hypersensitivity sites, split by previously reported signal strength (c), or CpG islands (d)^11^. Shaded area indicates the standard deviation across single cells. (**e**) Averaged single-cell DNA accessibility (blue) and DNA methylation (red) profiles at TSS, segregated by gene expression levels: High (solid), medium (dashed), low (dotted), or undetected (dash dot). (**f**) Boxplot of Pearson’s correlation between gene expression and indicated epigenetic features for individual cells (black) at *cis* regulatory elements. Blue dot indicates averaged single-cell correlation, and red square indicates pseudo-bulk correlation. Data in panels (b)-(f) are obtained by applying scMAT-seq to individual H9 hESCs. (**g**, **h**, **i**) H9 (purple) and HEK293T (orange) cells can be separated based on their transcriptome (g), DNA accessibility (h), or DNA methylation (i).

To successfully implement scMAT-seq, we optimized two important steps of this method. First, we evaluated different buffer conditions (1x first strand buffer, 1x GC buffer, or 50:50 ratio of each) to ensure that the simultaneous reverse transcription of mRNA and marking of open chromatin by M.CviPI is efficient. While the number of transcripts and the total number of methylated cytosines detected were similar across all buffer conditions (Supplementary Fig. 1a-c), we found that the ratio of exogenous to endogenous methylated cytosines was significantly lower in the 1x first strand buffer, indicating inhibition of M.CviPI in this buffer (Supplementary Fig. 1d). Therefore, we used a 50:50 buffer ratio for all further experiments (Supplementary Fig. 1e). Next, a consequence of making three different measurements from the same cell without physical separation of nucleic acids prior to amplification is that detection of the less abundant type of molecule requires sequencing the libraries to higher depths, thereby increasing the cost of the method. For example, in our initial implementation of scMAT-seq, only 4.2% of molecules in the library derived from mRNA, requiring higher sequencing depths for cell type identification (Supplementary Fig. 2a), a limitation we also observed in our previous single-cell multiomics measurements^10^. To overcome this, we developed an mRNA enrichment protocol after IVT where poly-A mRNA derived amplified RNA molecules were selectively separated from other molecules using biotinylated poly-A primers and streptavidin coated magnetic beads (Fig. 1a). To maximize mRNA enrichment, we tested four commercially available beads, and found that C1 and M270 beads provided the best enrichment, with a significant increase in the number of transcripts and genes detected (Supplementary Fig. 2b-d). Additionally, we found that library preparation could be performed directly on beads without impacting efficiency, thereby simplifying the enrichment protocol (Supplementary Fig. 2e,f). After enrichment, we observed a 9.4 fold increase in mRNA-derived molecules (40.0%), along with a significant increase in gene detection at levels comparable to other single-cell mRNA sequencing techniques (Fig. 1b and Supplementary Fig. 2a)^8^. Finally, the transcriptome obtained after enrichment was highly correlated (Pearson’s *r* = 0.95) to the non-enriched transcriptome, showing that the mRNA enrichment protocol does not introduce biases in quantifying gene expression (Supplementary Fig. 2g).

As proof-of-concept, we first applied scMAT-seq to H9 human embryonic stem cells (hESC). Comparison with previously published DNase I hypersensitivity sites (DHS) showed that, as expected, more accessible sites were associated with larger peaks and greater domain spreading in scMAT-seq (Fig. 1c)^11^. Similarly, results from scMAT-seq displayed a highly monotonic relationship with DHS scores (Spearman’s *p* = 0.99), showing that scMAT-seq faithfully reproduces a widely used technique for profiling DNA accessibility (Supplementary Fig. 3a). Furthermore, while DNA accessibility profiling is typically performed on fresh samples to maintain chromatin structure, we tested if scMAT-seq could be extended to frozen samples. When compared to freshly processed cells, we found that sorted cells stored at −80°C produced similar genome-wide profiles of DNA accessibility at DHS and transcription start sites (TSS), demonstrating that scMAT-seq can be applied to a larger spectrum of cryopreserved samples (Supplementary Fig. 4a-c). Next, we validated how accurately scMAT-seq captures the methylome of single cells. scMAT-seq builds upon a method we recently developed (scMspJI-seq), where we showed that scMspJI-seq is an alternate approach to single-cell bisulfite sequencing^12^. As with scMspJI-seq, we find that greater than 97% of the 5mC sites detected by scMAT-seq overlapped with published bulk bisulfite sequencing (Supplementary Fig. 3b)^13^. In addition, we observed global demethylation at CpG islands (CGI), consistent with hypomethylation at most CGIs within mammalian genomes (Fig. 1d)^14^. To verify that scMAT-seq can capture the relationship between DNA accessibility and DNA methylation, and its impact on gene expression, we segregated genes based on their expression level to find that increasing levels of gene expression are associated with more open chromatin and reduced DNA methylation at TSSs (Fig. 1e). Interestingly, when compared directly at the single-cell level, we observed higher correlations between gene expression and DNA accessibility that is computed over both the promoter and gene body instead of just the promoter alone (Fig. 1f). Further, as seen previously by Clark *et al*., the pseudo-bulk correlation between gene expression and DNA accessibility in the promoter region is higher than the average of individual cells (Pearson’s *r* of 0.80 and 0.28, respectively), likely due to the small size of the promoter region which limits the detection of reads in these regions in single cells^4^. Together, these results demonstrate that scMAT-seq can be used to quantify DNA accessibility, DNA methylation and the transcriptome from the same cell. Finally, to demonstrate that scMAT-seq can be used to identify distinct cell types and construct cell type-specific epigenetic landscapes, we performed scMAT-seq on HEK293T cells. As expected, RNA expression data from scMAT-seq could be used to easily distinguish between the two cell lines H9 and HEK293T (Fig. 1g). Similarly, the cell lines could be segregated by the first principal component for both DNA accessibility and DNA methylation, suggesting that scMAT-seq can successfully capture cell type-specific epigenetic profiles (Fig. 1h,i).

Next, we applied scMAT-seq to study early events in post-implantation human development. While epigenetic regulation of gene expression during post-implantation development and gastrulation have recently been characterized using a similar multiomics singlecell technique in mice, similar studies in human embryos have not been possible despite their fundamental importance to human health and disorders^2^. Further, morphological divergence from mice makes it challenging to directly extrapolate the emergence and maturation of different cell types during this period to human development. Therefore, to study human embryogenesis, we used an organoid model that mimics post-implantation amniotic sacs that can induce early germ layer lineages, similar to previous systems (Fig. 2a)^15–21^. As expected, analysis of the transcriptome from scMAT-seq identified epiblast-like cells (EPILC) expressing pluripotency markers POU5F1, NANOG, SOX2 and DPPA4, as well as amniotic ectoderm-like cells (AMLC) expressing TFAP2A, GATA3, HAND1, BMP4, and CDX2 (Fig. 2b,c). Further, EPILCs in these asymmetric cysts have previously been shown to undergo epithelial to mesenchymal transition, mimicking gastrulation and presenting features of posterior primitive streak-/mesoderm-like cells (PPSLC)^15,19^. In agreement with those observations, we found cells expressing markers of the posterior primitive steak, including T, EOMES, LHX1, MESP1, MESP2, GATA6, LEF1, SNAI1 and SNAI2, suggesting that PPSLCs are also derived in our system (Fig. 2b,c). Unlike the other organoid models previously developed, we unexpectedly found a small number of anterior primitive streak-/endoderm-like cells (APSLC) within the same organoid system, expressing high levels of NODAL, FOXA2, SOX17, OTX2, CXCR4, and even expressing genes related to the organizer cell fate including GSC and signaling inhibitors DKK1 and CER1 (Fig. 2b,c). The emergence of these APSLCs indicates an area of relatively high Activin/Nodal signaling and low BMP signaling within the organoid^22^. In human embryos, in addition to its role at the anterior primitive streak, SOX17 is a critical regulator of primordial germ cells (PGC) specification and unlike in mice, it is upregulated before the commonly used PGC marker PRDM1 (also known as BLIMP1)^23^. Interestingly, a subset of cells were found to express both SOX17 and PRDM1, together with other PGC markers, including NANOG, POU5F1, NANOS3, and TFAP2C (also known as AP2-gamma), indicating the emergence of human primordial germ cell-like cells (hPGCLC) in our system (Fig. 2b,c).

**Figure 2.**
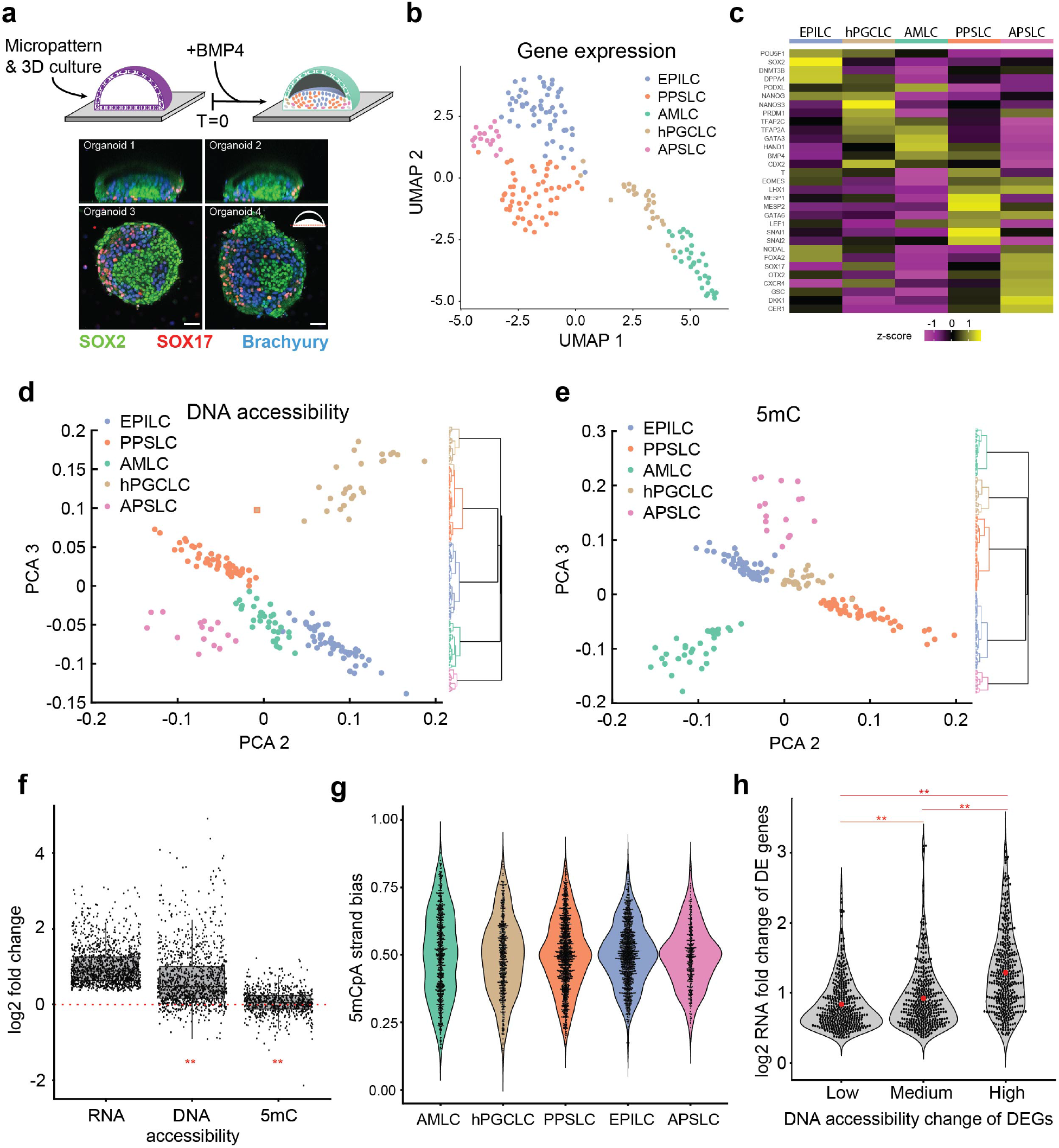
scMAT-seq maps the epigenome and transcriptome of cell types during human gastrulation using a 3D organoid model. (**a**) Schematic for the *in vitro* generation of 3D postimplantation amniotic sac organoids. Representative immunofluorescence staining of cross sections of the organoids 48 hours post BMP4 addition is shown for SOX2 (green), SOX17 (red), and Brachyury (blue). (**b**) UMAP visualization of the gastruloid based on the single-cell transcriptomes obtained from scMAT-seq. Cell types are assigned to clusters based on established marker genes. (**c**) Heatmap of z-scores for expression of marker genes for different cell types identified in the gastruloid. (**d**,**e**) Principal component projections show that cell types identified in the human gastruloids can also be distinguished using their DNA accessibility landscapes (d) or DNA methylation landscapes (e). Panels also show the corresponding dendrograms. A square border around dots indicate cell type identification by the transcriptome that differs from the epigenetic feature. (**f**) Boxplot of single-cell averaged log2 fold change in gene expression levels, promoter and gene body DNA accessibility levels, and gene body DNA methylation levels, for DEGs between two cell types. Dots indicate individual DEGs. Note that the log2 fold change in RNA is computed by taking the ratio of expression in a cell type where the gene is expressed at a higher level than in the other cell type. The log2 fold change in DNA accessibility and DNA methylation is shown for the same corresponding pair of cell types. ** indicates a statistically significant change (*p* < 0.01, based on bootstrapped distributions from non-differentially expressed genes) in the log2 fold change of the epigenetic features for DEGs relative to non-differentially expressed genes. (**g**) Violin plot of non-CpG methylation strand bias for each cell type. Strand bias is defined as the ratio of non-CpG methylated sites detected on the plus strand of a chromosome over all non-CpG methylated sites detected on a chromosome. Dots represent individual chromosomes in single cells. (**h**) Violin plot of log2 fold change in gene expression for DEGs partitioned by the relative change in DNA accessibility. Black dots indicate individual DEGs. Red dot indicates the mean of the distribution. ** indicates a statistically significant change (*p* < 0.01, two-sided Mann-Whitney U test) between the distributions.

In addition to discovering cell types corresponding to the early germ layer lineages and hPGCLCs, we discovered an unexpected population of cells with high expression of SOX2, PAX6 and PAX3, reminiscent of neuroectoderm cells, which typically arise later in development (Supplementary Fig. 5a,b)^24^. To investigate the origin of this population of cells, we performed a simpler experiment to test the differentiation potential of the starting induced pluripotent stem cells (iPSCs) by treating them with the GSK-3 inhibitor CHIR99021 for 24 hours (+CHIR), which is known to activate the WNT pathway, causing differentiation towards the mesodermal lineage in iPSCs^25^. iPSCs with or without CHIR treatment were probed with scMAT-seq, and clustering based on the transcriptome revealed 3 groups of cells – a pluripotent population within untreated cells expressing SOX2, POU5F1, NANOG, and DNMT3B, a mesodermal CHIR-treated population expressing T, EOMES, and NODAL, and a neuroectoderm-like cell (NELC) population present in both CHIR treated and untreated cells, expressing SOX2, PAX6, and PAX3, similar to that observed in the organoid (Supplementary Fig. 5c,d). These results indicate heterogeneity within the iPSC population, with a subset of cells that are biased towards the neuroectoderm lineage. Further, as this pre-existing population of NELCs was not responsive to CHIR treatment and displayed limited differentiation potential, it was removed from downstream analysis, highlighting the benefits of performing single-cell measurements on complex biological systems.

We next focused attention on the epigenomes of the different cell types identified in the gastruloid. Together with the rapid emergence of different cell types within a 48-hour window in the organoid, we found that the DNA accessibility and 5mC landscapes were reprogrammed, with distinct profiles for each cell type. Further, both genome-wide epigenetic features could be independently used to accurately cluster cells by cell type (Fig. 2d,e). Further, when comparing differentially expressed genes (DEGs) between different cell types, higher gene expression was generally associated with higher DNA accessibility (Fig. 2f and Supplementary Fig. 6a,b). In contrast, changes in gene expression were significantly correlated to changes in gene body DNA methylation in only a subset of cell types (Fig. 2f and Supplementary Fig. 6c). Together, these results indicate that reprogramming of DNA accessibility is a major driver of changes in gene expression and the emergence of distinct cell types during *in vitro* gastrulation, whereas gene body methylation plays a more limited role in tuning gene expression by regulating select genes and cell types. To gain better understanding of the dynamics of DNA methylation turnover during gastrulation, we observed that the expression level of the *de novo* DNA methyltransferase DNMT3B was lower in more differentiated cell types, especially PPSLC, PGCLC and AMLC, compared to EPILC, consistent with previous observations that DNMT3B is associated with pluripotency and is responsible for the high levels of non-CpG methylation in hESCs that drops substantially during differentiation (Fig. 2c and Supplementary Fig. 7a)^26^. In agreement with this, we observed that the levels of 5mCpA relative to 5mCpG dropped for PPSLC, PGCLC and AMLC compared to EPILC (Supplementary Fig. 7b). Further, similar to our recent work, scMAT-seq enables strand-specific quantification of DNA methylation, which provides insights into DNA methylation dynamics as increasing levels of asymmetric 5mCpA between the two strands of DNA implies a reduction in the rates of *de novo* methylation^6^. We found that PPSLC, PGCLC and AMLC showed greater strand-specific asymmetry in 5mCpA compared to EPILC, consistent with the expression levels of DNMT3B, suggesting that *in vitro* gastrulation is marked by a drop in genomewide non-CpG methylation and pluripotency, arising from a global reduction in the rates of *de novo* methylation (Fig. 2g and Supplementary Fig. 7c).

While changes in DNA accessibility and gene expression were generally correlated across all cell types, fold changes in the expression of individual DEGs did not always result in proportional fold changes in DNA accessibility (Fig. 2f and Supplementary Fig. 6a,b). To investigate this further, changes in DNA accessibility of DEGs were split into three groups, and as expected, larger increases in DNA accessibility were on average associated with larger increases in gene expression (Fig. 2h). However, interestingly we observed outlier genes in all groups that showed fold changes in expression that were significantly higher than the mean, with the group containing genes with the most open chromatin displaying a longer tail of highly DEGs compared to the other two groups. These results suggest that for a subset of genes, other epigenetic features potentially drive changes in gene expression that do not directly alter DNA accessibility within the promoter and gene body, highlighting that the combinatorial action of epigenetic regulators can non-additively tune gene expression.

Finally, we focused attention on the population of hPGCLCs that we identified 48 hours post BMP4 addition. Due to a lack of availability of human embryos at these stages of development, the precursors that specify human primordial germ cells (hPGCs) *in vivo* remain unclear^27,28^. In mice, it is well-established that mPGCs emerge from epiblast cells, and similar results have been found in the developing porcine embryo; however, in the nonhuman primate cynomolgus macaques, PGCs have been found to arise from the extra-embryonic dorsal amnion^29,30^. Further, there is significant divergence between human and mouse in the transcription factor network that is responsible for commitment towards the germ cell lineage, and therefore, the identity of the progenitors that specify hPGCs in humans remain unknown^27,28^. The hPGCLCs that we identified 48 hours post BMP4 addition expressed well-established markers of hPGCs, including SOX17, TFAP2C, PRDM1, NANOG, POU5F1, and NANOS3; however, they were found to not display any genome-wide erasure of DNA methylation, a characteristic feature of PGC maturation, suggesting these cells were pre-migratory PGCs that had recently undergone specification (Fig. 2c and Supplementary Fig. 7c)^23,28^. Therefore, to investigate the early events involved in germ cell specification, we systematically characterized younger organoids 20 and 36 hours post BMP4 addition. Using RNA expression data and Monocle 3, individual cells sequenced from the organoid were assigned a pseudotime to indicate the position of a cell along a differentiation trajectory (Fig. 3a)^31^. In general, we found the pseudotime assignment to correspond well with organoid age, providing confidence that it accurately described progress along a differentiation lineage (Fig. 3b). Similarly, as expected, one of the trajectories from EPILCs was found to bifurcate into two paths, developing into either PPSLCs or APSLCs (Fig. 3a). Strikingly, we found that the hPGCLCs and AMLCs bifurcate from a common progenitor population, which suggested the emergence of a precursor within 20 hours after treatment with BMP4. Compared to the EPILCs, we discovered that this progenitor population expressed genes related to the amnion (TFAP2A, GATA3 and CDX2) as well as gastrulating cells (EOMES and T), and downstream targets of BMP4 signaling (BAMBI, ID1-3 and MSX2) (Fig. 3c, Supplementary Fig. 8a and Supplementary Fig. 9). Notably, this was consistent with another recent observation in a disorganized 3D aggregates based system where hPGCLCs were found to emerge from a TFAP2A+ progenitor population^32^. Comparison with this system showed that the progenitors in our organoids are closest to day 1 cells post BMP4 addition in the disorganized aggregates, while hPGCLCs in our system are closest to day 2 cells in the disorganized aggregates (Fig. 3d). Further, we investigated DEGs characterizing the mesoderm, amnion and germ cells to find that these genes were more accessible and contained higher levels of gene body methylation in the progenitor population compared to EPILCs, suggesting that the progenitors are primed towards conversion to AMLCs and PGCLCs (Supplementary Fig. 8b-d). Overall, these results suggest that the progenitor population first emerges from EPILCs within 20 hours of BMP4 addition, with transient characteristics of both amniotic- and mesoderm-like cells, before getting specified towards hPGCLCs.

**Figure 3.**
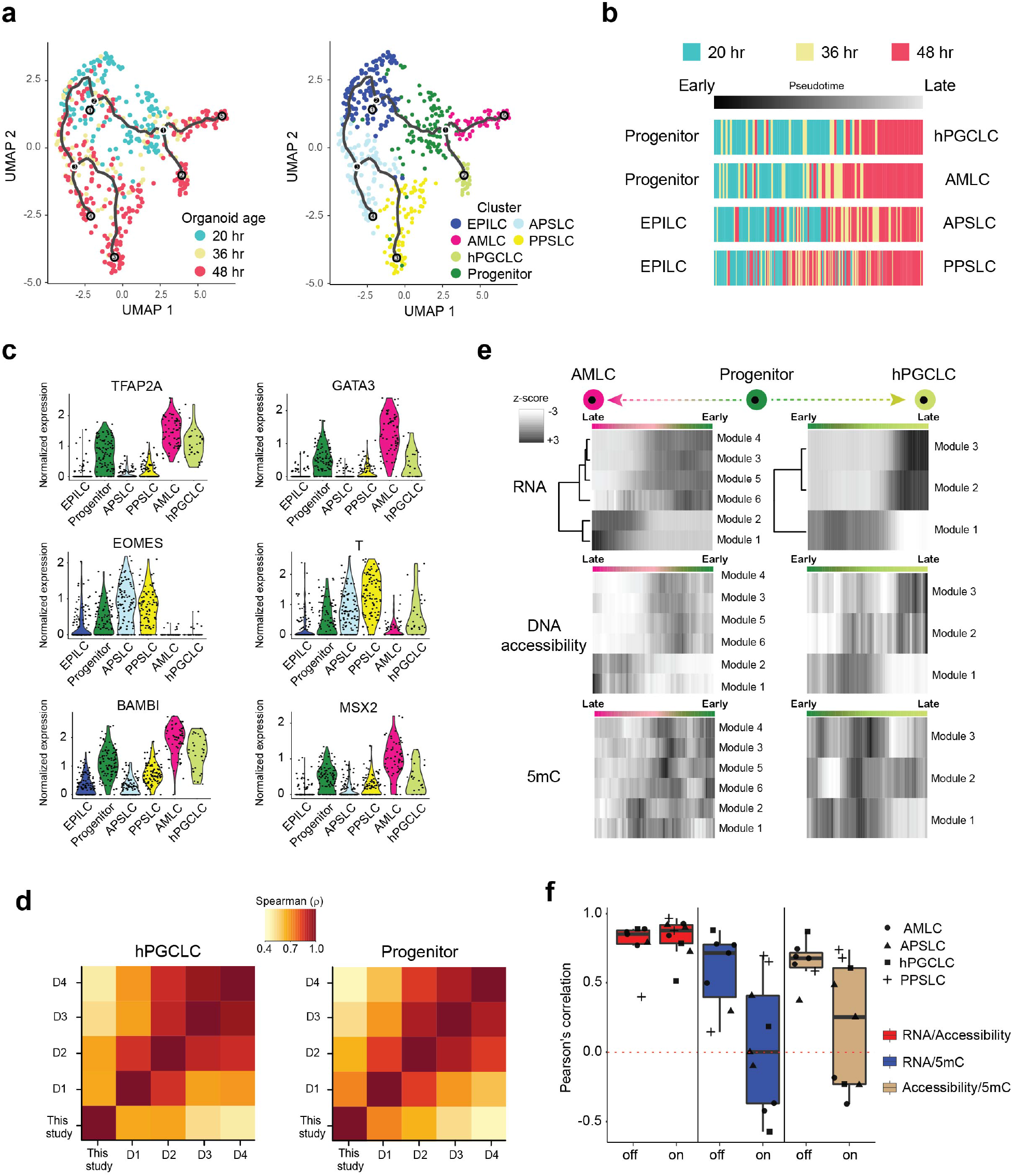
scMAT-seq characterization of time course experiment on gastruloids reveals the identity of hPGCLC progenitors. (**a**) (*Left*) UMAP visualization of the single-cell transcriptomes of the gastruloid at different timepoints, 20 hours (blue), 36 hours (tan), and 48 hours post BMP4 addition (ruby). The solid black line shows the inferred differentiation trajectory of cells from Monocle 3. (Right) Assignment of cell types to clusters identified in the time course experiment. (**b**) Relationship between predicted pseudotime trajectory and gastruloid age for different differentiation lineages. (**c**) Violin plots for select genes, highlighting that the progenitor population expresses markers of the amnion (TFAP2A and GATA3), gastrulating cell fate (EOMES and T), and downstream targets of BMP4 signaling (BAMBI and MSX2). (**d**) Heatmap of Spearman’s correlation coefficient comparing gene expression of hPGCLCs and progenitors identified in this study to that found in disorganized aggregates previously described by *Chen et al*.^32^. D1 to D4 indicates day 1 to day 4 post BMP4 addition in the disorganized aggregates. (**e**) Heatmap of z-scores for DEG-derived gene modules along the AMLC and hPGCLC pseudotime trajectory, together with the corresponding changes in DNA accessibility and DNA methylation. The color bar indicates position along the pseudotime, with the progenitor in green, AMLC in pink, and hPGCLC in light green. (**f**) Boxplot of Pearson’s correlation between the indicated variables along pseudotime with gene modules being divided into two groups – those that are upregulated (On) or those that are downregulated (Off) with pseudotime.

Finally, to systematically map changes in the epigenome along all differentiation trajectories during gastrulation, we grouped genes that varied throughout pseudotime into gene modules based on their expression similarity. In all trajectories, we found that changes in promoter and gene body DNA accessibility closely varied with changes in RNA trajectory (Fig. 3e and Supplementary Fig. 10). In contrast, gene body DNA methylation was associated to a lesser extent with the transcriptome (Fig. 3e and Supplementary Fig. 10). To quantify this relationship across all trajectories, gene modules were split into two groups – one that is downregulated (Off) and the other that is upregulated (On) through differentiation. We found that reprogramming of DNA accessibility was highly correlated to changes in gene expression for both genes turning Off and On (Fig. 3f). Surprisingly however, while gene body DNA methylation changes were generally well correlated to expression changes for genes that turn Off across all differentiation trajectories, we found a wide range of correlations, depending on the trajectory, for genes that turn On (Fig. 3f). Similarly, direct comparison of DNA accessibility and DNA methylation showed high correlation for genes that are downregulated while the correlation was lower, and trajectory dependent, for genes that are upregulated during differentiation (Fig. 3f). Previous work has shown that more open chromatin in highly expressed genes is linked with greater access for the *de novo* methylation machinery, resulting in higher gene body methylation^33^. We hypothesize that the high correlation between DNA accessibility and gene body DNA methylation for genes that are turning Off is possibly due to these regions being inaccessible and therefore, independent of the expression level of the *de novo* DNA methyltransferases. In contrast, for genes that are turning On, we observe a wide range of trajectory-dependent correlations, with differentiation towards AMLCs displaying lower correlation than APSLC and PPSLC lineages, possibly due to the lower expression of DNMT3B resulting in reduced *de novo* methylation activity within gene bodies of AMLCs (Fig. 3f, 2c and Supplementary Fig. 7a). Together, this suggests that increasing DNA accessibility and not gene body methylation is required for gene activation, and that while high DNA accessibility in gene bodies could lead to high DNA methylation, this correlation is decoupled by low *de novo* methyltransferase activity.

## Conclusions

In this report, we developed a multiomics single-cell method scMAT-seq to simultaneously quantify DNA accessibility, DNA methylation and mRNA from the same cell without requiring physical separation of the nucleic acids prior to amplification, enabling efficient and high-throughput mapping of cell types and their corresponding epigenetic profiles *in silico*. When applied to human gastruloids, we show that both the transcriptome and the epigenetic features can be used to identify different cell types. Notably, we discovered that the progenitor population that gives rise to hPGCLCs emerge from EPILCs with transient epigenetic and transcriptional signatures of both amnion- and mesoderm-like cells. In summary, scMAT-seq provides an integrative approach to investigate the role of epigenetic features in regulating gene expression and cell fate decisions in complex and dynamic biological systems.

## Methods

### Mammalian cell culture

All mammalian cells were maintained in incubators at 37°C and 5% CO_2_. HEK293T cells were cultured on tissue culture treated plastic in high glucose DMEM (Gibco, 10569044) containing L-Glutamine and sodium pyruvate, supplemented with 10% FBS (Gibco, 10437028) and 1x Penicillin-Streptomycin (Gibco, 15140122). H9 cells and hiPSCs (Allen Cell Collection, line AICS-0024) were grown feeder-free on Matrigel (Fisher Scientific, 08-774-552) coated plates in mTeSR1 medium (STEMCELL Tech., 85850). Cells were routinely passaged 1:6 once they reached 75% confluency using 0.25% trypsin-EDTA (Gibco, 25200056) for HEK293T cells, Versene solution (Gibco, 15040066) for H9 cells, and ReLeSR (STEMCELL Tech., 100-0484) for hiPSCs. For FACS sorting, a single-cell suspension was made using 0.25% trypsin-EDTA. The trypsin was then inactivated using serum containing medium. Afterwards, the cells were washed with 1x PBS before being passed through a cell strainer and sorted for single cells into 384-well plates. Use of the H9 human embryonic stem cell line was approved by the Human Stem Cell Research Oversight Committee (Protocol: hSCRO# 2019-1084).

### hiPSC derived mesoderm cell culture

Mesoderm differentiation was performed on hiPSCs (Allen Cell Collection, line AICS-0024). For mesodermal differentiation, the media was replaced with mTeSR1 supplemented with 5 uM of CHIR99021 (STEMCELL Tech., 72052) for 24 hours. Afterwards, the cells were dissociated into a single cell suspension using TrypLE reagent (Gibco, 12563011), and gentle pipetting. TrypLE was then inactivated using a serum containing medium. Next, the cells were washed with 1x PBS before being passed through a cell strainer and sorted for single cells into 384-well plates.

### Post-implantation amniotic sac organoid culture

The development of micropatterned 3D stem-cell cultures with a single lumen is described in Karzbrun *et al*.^21^. We applied the protocol here as follows:

#### Microfabrication of PDMS stamps

PDMS stamps were created with circular features 250 μm in diameter. Stamps were prepared using standard soft-lithography techniques on a four-inch wafer. One layer of photoresist (Microchem, SU-8 2075) is spun onto a silicon wafer at a thickness of 100 μm. Photoresist is exposed to ultraviolet light using a mask aligner (Suss MicroTec, MA6) and unexposed photoresist is developed away to yield multiple arrays of posts. A trimethylchlorosilane layer is vapor deposited on the developed wafer to prevent adhesion. A 10:1 ratio of PDMS and its curing agent (Dow Corning, SYLGARD 184 A/B) is poured onto the wafers and cured at 65°C overnight. The PDMS layer is then peeled off the silicon mold and individual stamps are cut out using a razor blade for future use.

#### Micro-contact printing

Sterile PDMS stamps and 35 mm diameter custom-made glass-bottomed culture dishes are plasma treated for 1 minute on high setting (Harrick Plasma, PDC-32G) to activate both surfaces. Stamps are pressed features-side to the glass surface and held in place. To passivate the glass surface in nonpatterned regions, 0.1 mg/mL PLL-g-PEG solution (SuSoS AG, Switzerland) is added to the petri dish immediately after securing stamps to the glass surface and incubated for 30 minutes. Stamps are then carefully removed and stamped glass dishes are rinsed several times with PBS containing calcium and magnesium (PBS++). Laminin-521 (STEMCELL Tech., 77003) is added at a dilution of 5 μg/mL in PBS++ to incubate overnight at 4°C. The following day, stamped glass dishes are rinsed with PBS++ to remove excess unbound, laminin and used within 1-2 weeks.

#### Stem-cell seeding, lumen formation and differentiation

On Day 1: hiPSCs (Allen Cell Collection, line AICS-0024) are released from well-plate surfaces using non-enzymatic agitation following manufacturer’s instructions (ReleSR, STEMCELL Tech.). Cells are resuspended as a single-cell suspension at densities of 750K-1M cells/mL in mTeSR1 containing 10 μM ROCK inhibitor Y27632 (Abcam, ab120129). 200 μL of cell suspension is then pipetted onto prepatterned dishes and allowed to settle for 15 minutes before adding 1 mL of mTeSR1 and allowing cells to settle for 10 additional minutes. Excess media is aspirated, leaving enough liquid to cover patterns and is replaced with fresh 2 mL of mTeSR1. On day 2: mTESR1 media is exchanged with mTESR1 media containing Matrigel (4%, v/v). This triggers lumen formation over 24 hours. On Day 3: Micropatterned colonies have formed a lumen. mTESR1 is exchanged with the addition of 5 ng/ml Recombinant Human BMP4 (Fisher Scientific, 314BP010). Exposure to BMP4 triggers differentiation of cells and is considered 0 hours for experimental purposes. On Day 4 or 5: Samples were collected for single-cell sequencing at either 20, 36, or 48 hours post BMP4 supplementation. Samples were dissociated into a single-cell suspension using TrypLE, and gentle pipetting. TrypLE was then neutralized using a serum containing medium. Next, the cells were washed with 1x PBS before being passed through a cell strainer. Finally, single cells were sorted into 384-well plates using FACS.

### scMAT-seq

4 μL of Vapor-Lock (QIAGEN, 981611) was manually dispensed into each well of a 384-well plate using a 12-channel pipette. All downstream dispensing into 384-well plates were performed using the Nanodrop II liquid handling robot (BioNex Solutions). To each well, 100 nL of uniquely barcoded 7.5 ng/μL reverse transcription primers containing 4 nucleotide unique molecule identifiers (UMI) was added. The reverse transcription primers used here were previously described in Grun *et al*.^34^. Next, 100 nL of lysis buffer (0.175% IGEPAL CA-630, 1.75 mM dNTPs, 1:1,250,000 ERCC RNA spike-in mix (Ambion, 4456740), and 0.19 U RNase inhibitor (Clontech, 2313A)) was added to each well. Single cells were sorted into individual wells of a 384-well plate using FACS. Unless cryopreserved, after sorting, plates were heated to 65°C for 3 minutes and returned to ice. Next, 150 nL of reverse transcription GC tagging mix (0.7 U RNAseOUT (Invitrogen, 10777-019), 1.17x first strand buffer, 11.67 mM DTT, 3.5 U Superscript II (Invitrogen, 18064-071), 0.19 mM SAM, 1.17x GC reaction buffer, 0.1 U M.CviPI (NEB, M0227S)) was added to each well and the plates were incubated at 37°C for 1 hour, 4°C for 5 min, 65°C for 10 min, and 70°C for 10 min. Thereafter, 1.75 μL of second strand synthesis mix (1.74x second strand buffer (Invitrogen, 10812-014), 0.35 mM dNTP, 0.14 U E.coli DNA Ligase (Invitrogen, 18052-019), 0.56 U E.coli DNA Polymerase I (Invitrogen, 18010-025), 0.03 U RNase H (Invitrogen, 18021-071)) was added to each well and the plates were incubated at 16°C for 2 hours. Following this step, 400 nL of protease mix (6 μg protease (Qiagen, 19155), 6.25x NEBuffer 4 (NEB, B7004S)) was added to each well, and the plates were heated to 50°C for 15 hours, 75°C for 20 minutes, and 80°C for 5 minutes. Next, 500 nL of 5hmC-blocking mix (1 U T4-BGT (NEB, M0357L), 6x UDP-glucose, 1x NEBuffer 4) was added to each well and the plates were incubated at 37°C for 16 hours. Afterwards, 500 nL of protease mix (2 μg protease, 1x NEBuffer 4) was added to each well, and the plates were heated to 50°C for 3 hours, 75°C for 20 minutes, and 80°C for 5 minutes. Next, 500 nL of MspJI digestion mix (1x NEBuffer 4, 8x enzyme activator solution, 0.1 U MspJI (NEB, R0661L)) was added to each well and the plates were incubated at 37°C for 4.5 hours, and 65°C for 25 minutes. Unless otherwise noted, to each well, 320 nL of uniquely barcoded 125 nM double-stranded adapters were added. The double-stranded adapters have previously been described in Sen *et al*.^6^. Next, 680 nL of ligation mix (1.47x T4 ligase reaction buffer, 5.88 mM ATP (NEB, P0756L), 140 U T4 DNA ligase (NEB, M0202M)) was added to each well, and the plates were incubated at 16°C for 16 hours. After ligation, reaction wells receiving different barcodes were pooled using a multichannel pipette, and the oil phase was discarded. The aqueous phase was then incubated for 30 minutes with 1x AMPure XP beads (Beckman Coulter, A63881), placed on a magnetic stand and washed twice with 80% ethanol before eluting the DNA in 30 μL of nuclease-free water. After vacuum concentrating the elute to 6.4 μL, library preparation was performed as previously described in the scAba-seq and scMspJI-seq protocols^6,7^. Libraries were sequenced on an Illumina NextSeq 500 or an Illumina Hiseq 4000, sequencing a minimum of 75 bp on read 1 to detect methylated cytosines. A minimum of 25 bp on read 1 and 50 bp on read 2 was used to detect mRNA. Additionally, unless otherwise stated, detection of mRNA was only performed on RNA enriched samples and detection of methylated cytosines was performed only on unenriched samples.

### Optimizing buffer for simultaneous reverse transcription and GpC methylation tagging

In all experiments, 200 nL of the reverse transcription GC tagging mix was used and the 384-well plate was incubated at 37°C for 1 hour, 4°C for 5 minutes, 65°C for 10 minutes, and 70°C for 10 minutes. For the first strand buffer experiment, this reverse transcription GC tagging mix consisted of 0.7 U RNAseOUT, 2x first strand buffer, 20 mM DTT, 3.5 U Superscript II, 0.16 mM SAM, and 0.1 U M.CviPI. For the GC buffer experiment, this mix consisted of 0.7 U RNAseOUT, 3.5 U Superscript II, 0.16 mM SAM, 2x GC reaction buffer, 0.1 U M.CviPI, and 6 mM of MgCl_2_. For the 50:50 experiment, this reverse transcription GC tagging mix consisted of 0.7 U RNAseOUT, 1x first strand buffer, 1x GC reaction buffer, 10 mM DTT, 3.5 U Superscript II, 0.16 mM SAM, and 0.1 U M.CviPI. Following this, all other library construction steps were similar to the optimized scMAT-seq procedure. The volume of some reactions were altered as follows. Second strand synthesis was performed by adding 1.3 μL of second strand synthesis mix. The initial protease step was performed by adding 300 nL of protease mix. No 5hmC-blocking mix or secondary protease mix was added. After MspJI digestion, 200 nL of uniquely barcoded 1 nM or 200 nM double-stranded adapters were added. Afterwards ligation was performed by adding 800 nL of ligation mix. All other steps of library construction were unchanged.

### RNA enrichment

After *in vitro* transcription, 6 μL of amplified RNA product (aRNA) was combined with 2 μL of 1 μM biotinylated polyA primer (Integrated DNA Technologies, standard desalting, 5’-AAAAAAAAAAAAAAAAAAAAAAAA/3BioTEG/ -3’) and incubated for 10 minutes at room temperature. During this incubation, Dynabeads MyOne Streptavidin C1 beads (Invitrogen, 65001) were made RNase-free following the directions of the manufacturer. In addition, 2x and 1x B&W solution was made according to the manufacturer’s directions. After establishing RNase-free conditions, the beads were resuspended in 8 μL of 2x B&W solution. After the 10-minute incubation of aRNA with the biotinylated polyA primer, the beads were mixed in with the solution and incubated for 15 minutes at room temperature with constant shaking at 300 rpm using a thermomixer. Using a magnetic stand, the beads were separated from the supernatant, and the supernatant was discarded. The beads were washed twice with 1x B&W solution. After washing, the beads were resuspended in 10 μL of nuclease-free water. For on-bead processing, this product was taken to reverse transcription. For heat denatured processing, this product was heated to 70°C for 2 minutes, then a strong magnet was used to quickly separate the beads from the supernatant, and the supernatant was transferred to a new tube. In both cases, 5 μL of RNA enriched product was used for reverse transcription, and following this step, library preparation was performed as previously described in the scAba-seq and scMspJI-seq protocols^6,7^. For experiments using other bead types, the C1 beads were exchanged for other streptavidin beads, M270, M280, or T1 (Invitrogen, 65801D), with no other changes.

### scMAT-seq analysis pipeline

The scMAT-seq analysis pipeline was performed as described previously in Sen *et al*. with minor adjustments^6^. The custom Perl script used to identify 5mC positions in the genome was modified to interrogate the base prior to the called 5mC mark. Called 5mC marks preceded by a G were assigned to the DNA accessibility dataset, and due to the potential off target activity of M.CviPI, only those preceded by an A or T were assigned as endogenous 5mC marks, that were further split by their context (5mCpG, 5mCpA, 5mCpC, or 5mCpT). Custom codes for analyzing scMAT-seq data and the accompanying documentation is provided with this work (Supplementary Software).

The transcriptome analysis pipeline was similar to that described previously in Grün *et al*. with the following minor adjustments^34^. The right mate of paired-end reads was mapped in the sense direction using BWA (version 0.7.15-r1140) to the RefSeq gene model based on the human genome release hg19, with the addition of the set of 92 ERCC spike-in molecules. Any read mapping to multiple loci were distributed uniformly across those loci. Gene isoforms were consolidated into a gene count, and the UMIs were used to deduplicate reads and provide singlemolecule transcript counts for each gene in individual cells^34^. Genes that were not detected in at least one cell were removed from any downstream analysis.

### Comparison of scMAT-seq to established techniques

DNase I hypersensitivity sites for H9 and HEK293T cells were downloaded from UCSC table browser and sites were grouped based on detection scores^11^. Additionally, the genomic coordinates of CpG islands were downloaded from the UCSC table browser. To compare across datasets, the data was normalized to counts per million and a 75 base pair moving average was plotted for each region of interest. When comparing differing genomic regions within a sample, each region was further normalized by methylated cytosines that were detected when MspJI-seq was performed on bulk H9 gDNA that had been GpC methylated after stripping off chromatin. To do this, H9 gDNA was isolated using the DNeasy Blood & Tissue Kit (Qiagen, 69504). To 1.2 μg of purified H9 gDNA, 10 μL of protease mix (100 μg protease (Qiagen, 19155), 1x GC reaction buffer) was added, and the sample was heated to 50°C for 15 hours, 75°C for 20 minutes, and 80°C for 5 minutes. Next, 10 μL of GC tagging mix (8 U M.CviPI, 640 μM SAM, 1x GC reaction buffer) was added, and the sample was incubated at 37°C for 4 hours. Immediately after, 10 μL of additional GC tagging mix (4 U M.CviPI, 960 μM SAM, 1x GC reaction buffer) was added, and the sample was incubated at 37°C for 4 hours and 65°C for 20 minutes. Afterwards, 100 ng of this DNA was directly used as input to a scaled-up version of the scMspJI-seq protocol^6^.

5mC sites detected by scMAT-seq in H9 cells were compared to bulk bisulfite sequencing (GSM706061)^13^. A site was considered methylated if any level of 5mC was detected in the bulk bisulfite or pseudo-bulked scMAT-seq sequencing data. Overlapping 5mCpG sites were counted and compared to the number of non-overlapping sites.

### Genome-wide DNA accessibility and 5mC analysis in scMAT-seq

DNA accessibility and 5mC were quantified within 5 kb bins and then converted to binary scores. Further, pseudobulk profiles were generated using the assigned cell type from the transcriptome. For comparison between H9 and HEK293T cell lines, the top 2% most variable bins between groups were retained. For comparison in the post-implantation amniotic sac organoid, the top 1% most variable bins between groups were retained. After removal of bins with low variance, principal component analysis was performed on the remaining bins for individual cells, and hierarchical clustering was used to assign clusters. Cluster identification was performed through comparison to transcriptome derived cell types, where high similarity was observed between cluster calling for all 3 measurements.

### Promoter and gene body DNA accessibility and gene body 5mC analysis in scMAT-seq

For the quantification of DNA accessibility and 5mC in individual cells, a small pseudo-count was added prior to estimating the number of UMIs per million counts for each gene. The promoter of a gene was considered as 2,000 base pairs upstream of the transcription start site. Genes with low detection in all cells were removed from downstream analysis for that epigenetic feature.

### Gene expression analysis

The standard analysis pipeline in Seurat (version 3.1.5) was used for single-cell RNA expression normalization and analysis^35^. For H9 and HEK293T cell lines, cells containing more than 1,000 genes and more than 1,000 unique transcripts, as well as less than 20% ERCC spike-ins, were used for downstream analysis. For the post-implantation amniotic sac organoid, cells containing more than 1,000 genes and more than 4,000 unique transcripts, as well as less than 20% ERCC spike-ins, were used for downstream analysis. The default NormalizeData function was used to log normalize the data. In post-implantation amniotic sac organoids, the cluster identified as NELCs was removed from downstream analysis except where otherwise stated. When analyzing the time course data from the organoid, the FindIntegrationAnchors and IntegrateData functions were used to remove batch- and technique-specific effects. Thereafter, principal components were obtained from the 2,000 most variable genes and the elbow method was used to determine the optimal number of principle components used in clustering. UMAP based clustering was performed by running the following functions, FindNeighbors, FindClusters, and RunUMAP. After clustering, cell types were assigned to groups using known expression markers. To identify DEGs, the FindAllMarkers or FindMarkers function was used. The Wilcoxon rank sum test was used to classify a gene as differentially expressed, requiring a natural log fold change of at least 0.25 and an adjusted p-value of less than 0.05.

### Pseudotime analysis

UMAP coordinates and normalized gene expression data for highly variable genes was imported from Seurat to Monocle3 (version 0.2.1.5)^31^. Trajectories were built using the learn_graph function. Cells involved in the four observed trajectories were isolated separately using the choose_graph_segments function. Due to the bifurcation seen in the trajectories, some cells appeared in more than one of the trajectories. For each trajectory, the roots of the trajectories were chosen using the order_cells function to best correspond with cells from the 20-hour postimplantation amniotic sac organoids. After assigning a pseudotime to each cell, the genes varying over each pseudotime were determined using the graph_test function, with genes with a q-value under 0.01 and a Moran’s I value above 0.15 considered significant. Significantly varying genes for each trajectory were grouped into gene modules using the find_gene_modules function using a resolution value of 0.05. z-scores for gene expression of each gene module was calculated using the aggregate_gene_expression function. The corresponding z-score for DNA accessibility and 5mC was found and a 10-cell moving average was computed based on the pseudotime. For averaging and plotting, only cells passing quality controls for RNA expression, DNA accessibility and 5mC were considered.

## Acknowledgments

We thank Dr. Amander Clark at UCLA and members of the Dey lab for helpful discussions. We would like to thank Jennifer Smith at the Biological Nanostructures Laboratory in the California NanoSystems Institute (CNSI), supported by UCSB and UC Office of the President, for help with Illumina sequencing. Computational work was supported by the Center for Scientific Computing at CNSI and Materials Research Laboratory (MRL) at UCSB: an NSF MRSEC (DMR-1720256) and NSF CNS-1725797. A.H.K. was supported by the NSF Graduate Research Fellowship Program (Grant 1650114), and S.J.S was supported by NIH R21 HD099598-0. This work was supported by the NIH grant R01HG011013 to S.S.D.

## Author contributions

Conceptualization, A.C. and S.S.D.; Methodology, A.C. and S.S.D; Investigation, A.C., E.K. and A.H.K; Formal Analysis, A.C.; Writing – Original Draft, A.C.; Writing – Review & Editing, A.C. and S.S.D; Funding Acquisition, S.S.D.; Resources, M.J.R, S.J.S and S.S.D; Supervision, S.S.D.

