## Supplementary Information for "Integrated single-cell sequencing reveals principles of epigenetic regulation of human gastrulation and germ cell development in a 3D organoid model"

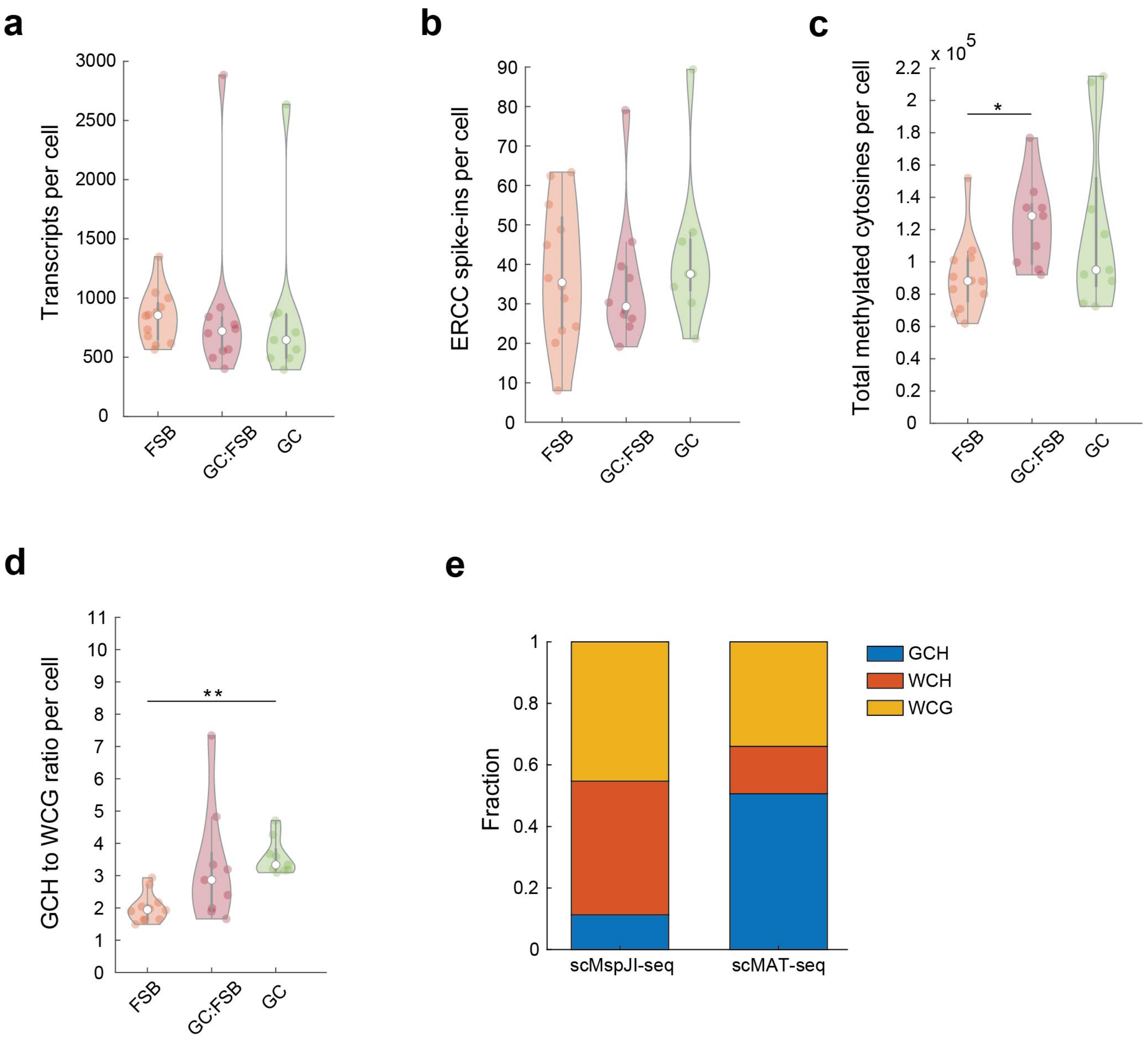


**Supplementary Figure 1 | Optimization of buffer conditions in scMAT-seq.** (**a**) Violin plot of the number of transcripts detected per cell in each buffer condition. (**b**) Violin plot of the number of synthetic ERCC RNA spike-in molecules detected in individual cells for different buffer conditions. (**c**) Violin plot of the total number of methylated cytosines detected, including both endogenous and exogenously introduced DNA methylation, in individual cells for different buffer conditions. (**d**) Violin plot of the ratio of methylated cytosines detected in different sequence contexts, comparing DNA accessibility (GCH) to endogenous CpG methylation marks (WCG), in individual cells for different buffer conditions. (**e**) Bar plot comparing detection of methylated cytosines in scMspJI-seq and scMAT-seq using optimized buffer conditions. The relative fractions of DNA accessibility (GCH), endogenous CpG methylation (WCG), and endogenous non-CpG methylation (WCH) are shown in blue, yellow, and orange, respectively. * and ** indicate $p < 0.05$ and $p < 0.01$, respectively (two-sided Mann-Whitney U test, Bonferroni-corrected p-values).

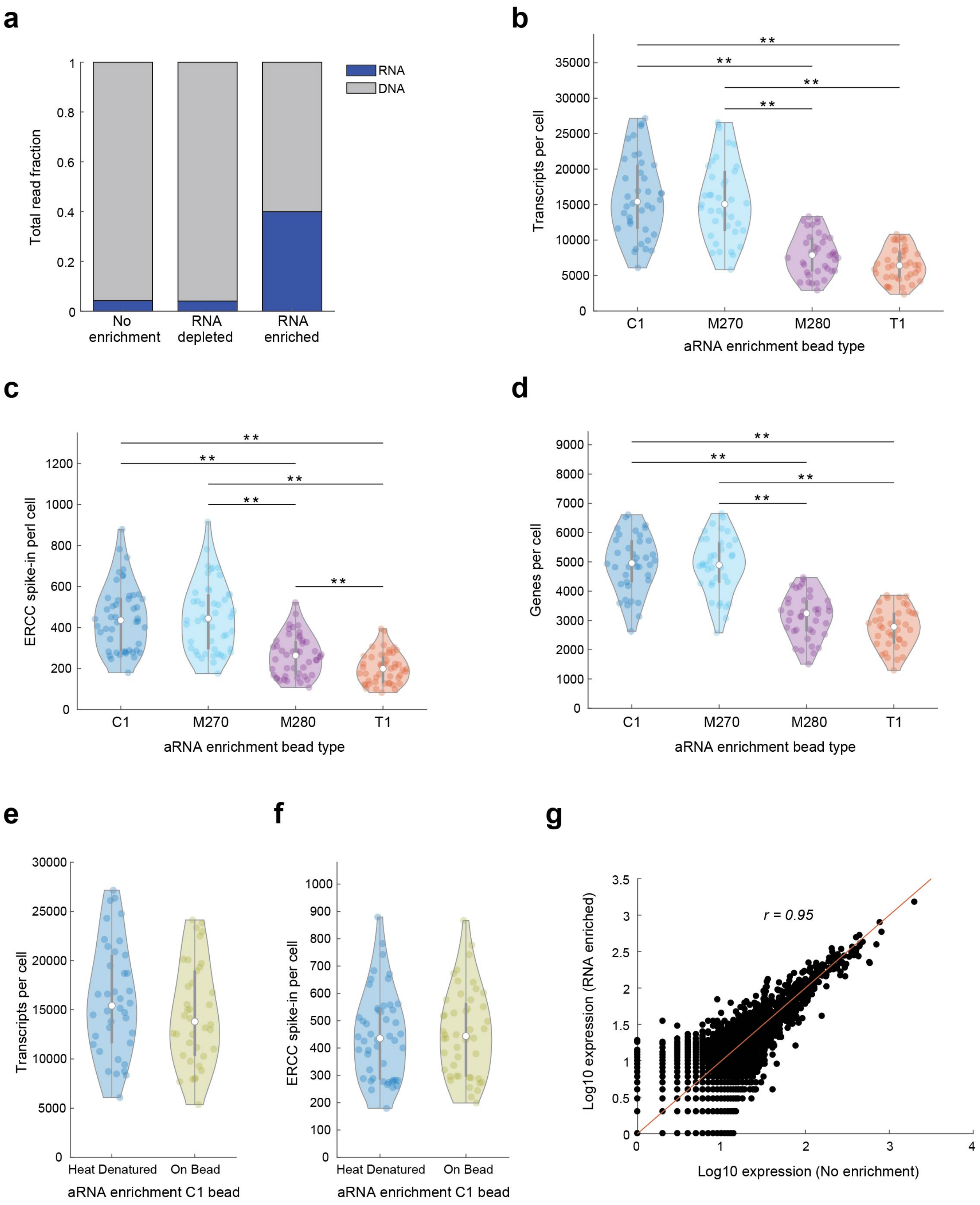


**Supplementary Figure 2 | Optimization of mRNA enrichment in scMAT-seq.** (**a**) Bar plot of the relative fraction of mRNA- and gDNA-derived reads in scMAT-seq without mRNA enrichment (No enrichment), with mRNA enrichment (RNA enriched), or the remaining flowthrough after mRNA enrichment (RNA depleted). (**b-d**) Violin plots of the number of transcripts (b), ERCC spike-in molecules (c), and genes (d) detected per cell using different magnetic streptavidin coated beads for mRNA enrichment. (**e**,**f**) Violin plots of the number of transcripts (e) and ERCC spike-in molecules (f) detected per cell after dissociating (heat denatured) or retaining (on bead) the aRNA on the magnetic streptavidin C1 beads. (**g**) Scatterplot comparing single-cell averaged gene expression with and without RNA enrichment from the same scMAT-seq sample. * and ** indicate $p < 0.05$ and $p < 0.01$, respectively (two-sided Mann-Whitney U test, Bonferroni-corrected p-values).

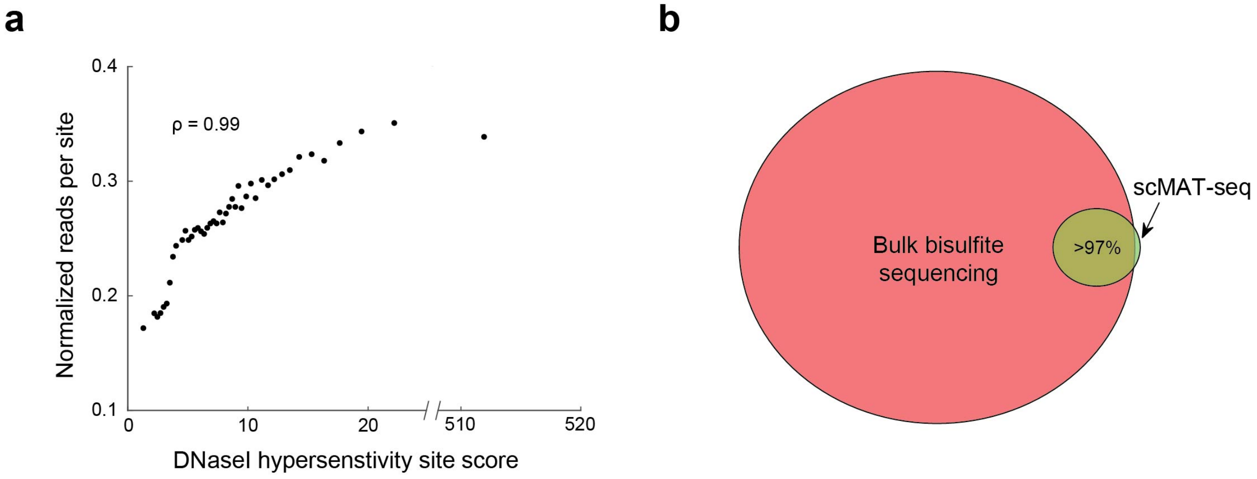


**Supplementary Figure 3 | scMAT-seq successfully reproduces DNA accessibility and DNA methylation profiles in hESCs.** (**a**) Scatterplot shows that single-cell averaged DNA accessibility in scMAT-seq is highly correlated to DNase I hypersensitivity scores (Spearman's $p = 0.99$)[1]. (**b**) Pie chart shows that greater than 97% of DNA methylation sites detected in single cells using scMAT-seq (green) are also detected in bulk bisulfite sequencing (pink)[2].

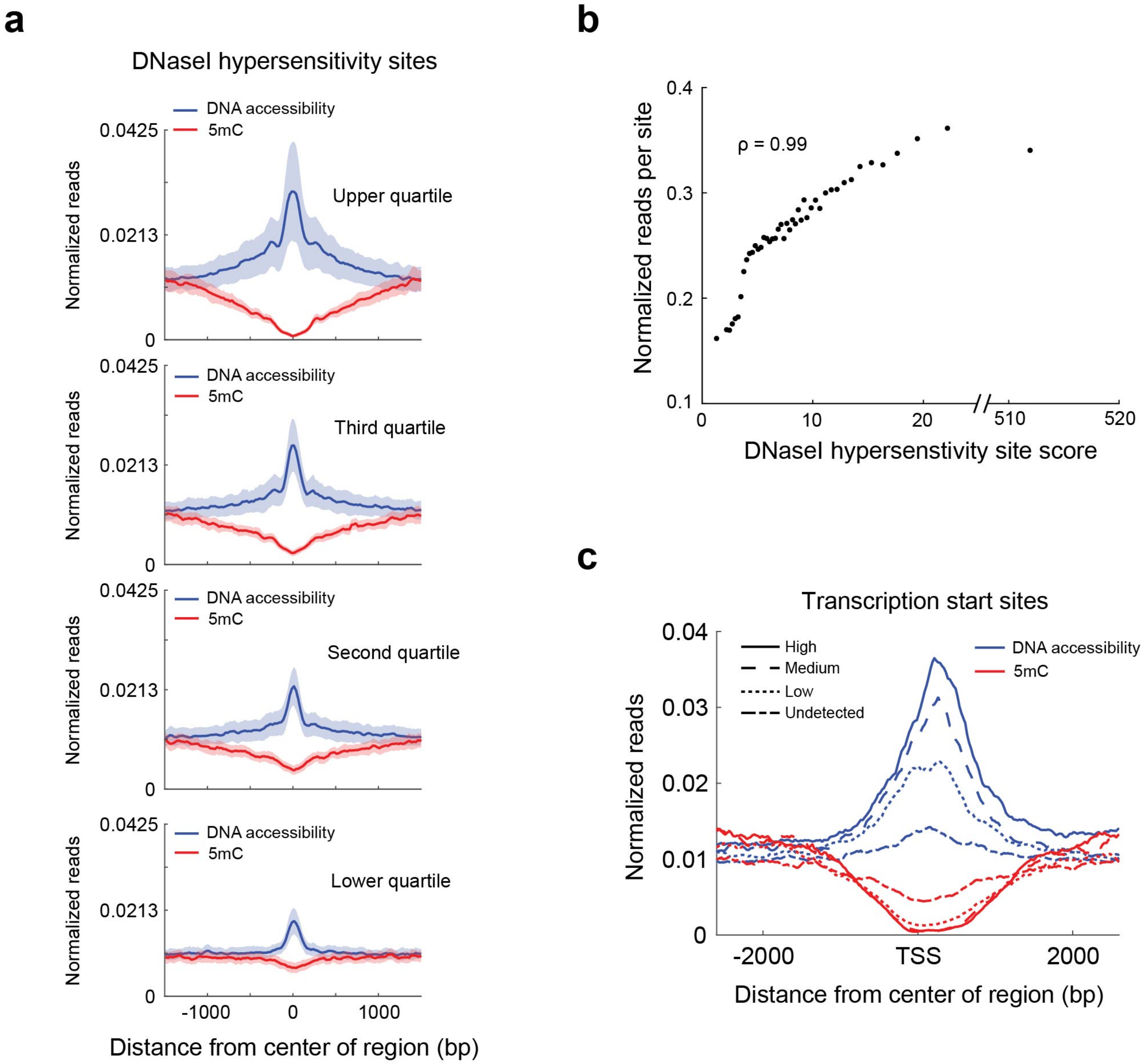


**Supplementary Figure 4 | scMAT-seq can accurately capture the genome-wide DNA accessibility and DNA methylation landscapes after cryopreservation of sorted samples.** (**a**) Averaged single-cell DNA accessibility (blue) and DNA methylation (red) profiles from cryopreserved hESCs at DNase I hypersensitivity sites, split by previously reported signal strength. Shaded areas indicate standard deviation across single cells[1]. (**b**) Scatterplot shows that averaged single-cell DNA accessibility in scMAT-seq from cryopreserved hESCs is highly correlated to DNase I hypersensitivity scores (Spearman's $p$ = 0.99)[1]. (**c**) Averaged single-cell DNA accessibility (blue) and DNA methylation (red) profiles in cryopreserved hESCs at TSS, segregated by gene expression levels: High (solid), medium (dashed), low (dotted), or undetected (dash dot).

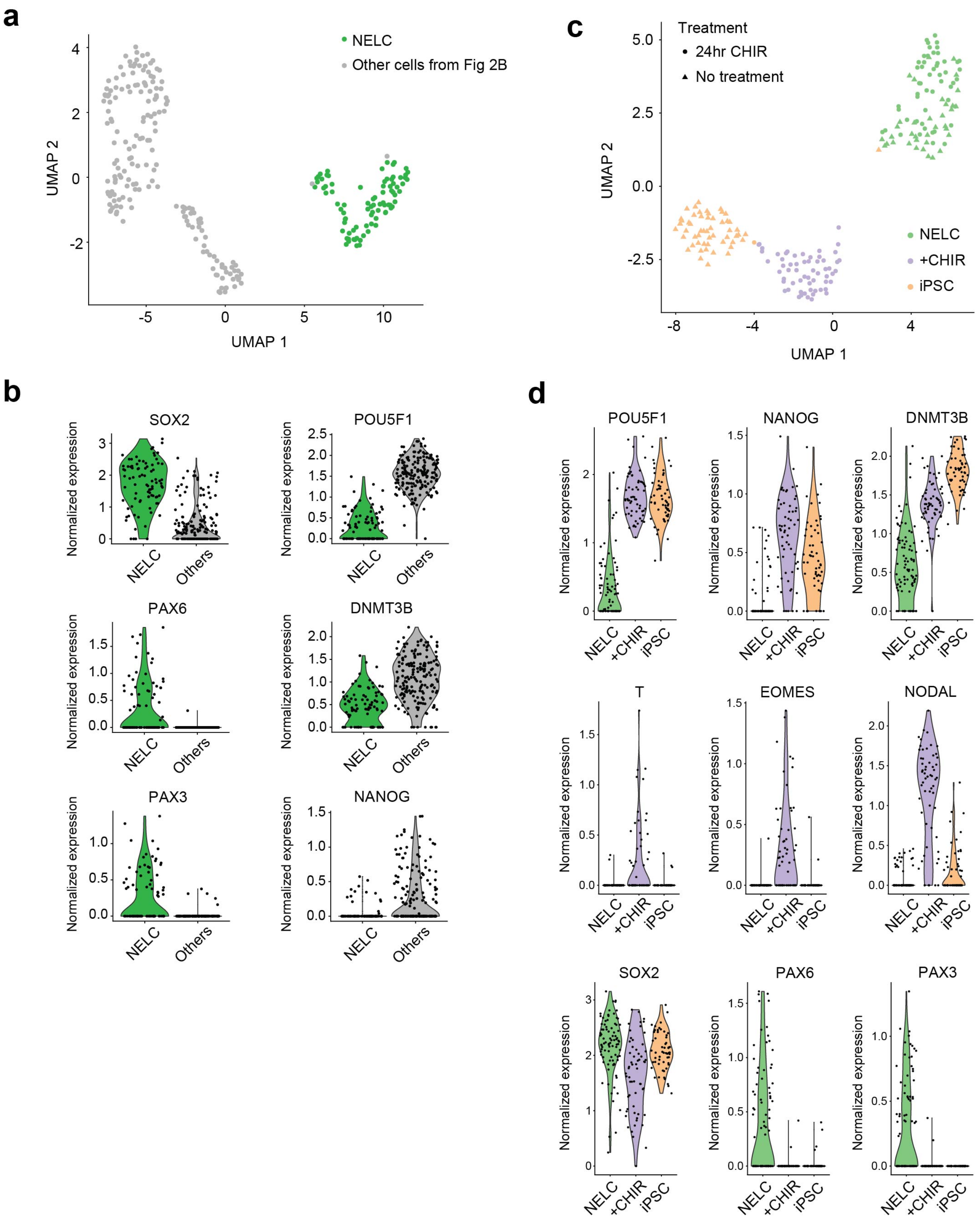


**Supplementary Figure 5 | Characterizing neuroectoderm-like cells during differentiation of iPSCs.** (**a**) UMAP visualization of single cells in the gastruloid 48-hour post BMP4 addition. Based on established marker genes, a cluster resembling NELCs is detected. (**b**) For the gastruloid characterized in (a), violin plots for select genes are shown, highlighting high expression of neuroectoderm genes (SOX2, PAX6, and PAX3) and low expression of pluripotency genes (POU5F1, DNMT3B, and NANOG) within the NELC population when compared to all other cells in the organoid. (**c**) UMAP visualization of untreated iPSCs (no treatment) and cells after treatment of iPSCs with CHIR99021 for 24-hours (24hr CHIR). Three populations of cells were detected, NELCs, iPSCs, and +CHIR cells. (**d**) For the system depicted in (c), violin plots for select genes are shown, highlighting high expression of pluripotency genes (POU5F1, DNMT3B, and NANOG) in iPSCs, high expression of mesodermal genes (T, EOMES, and NODAL) in the +CHIR cells, and high expression of neuroectoderm genes (SOX2, PAX6, and PAX3) in NELCs.

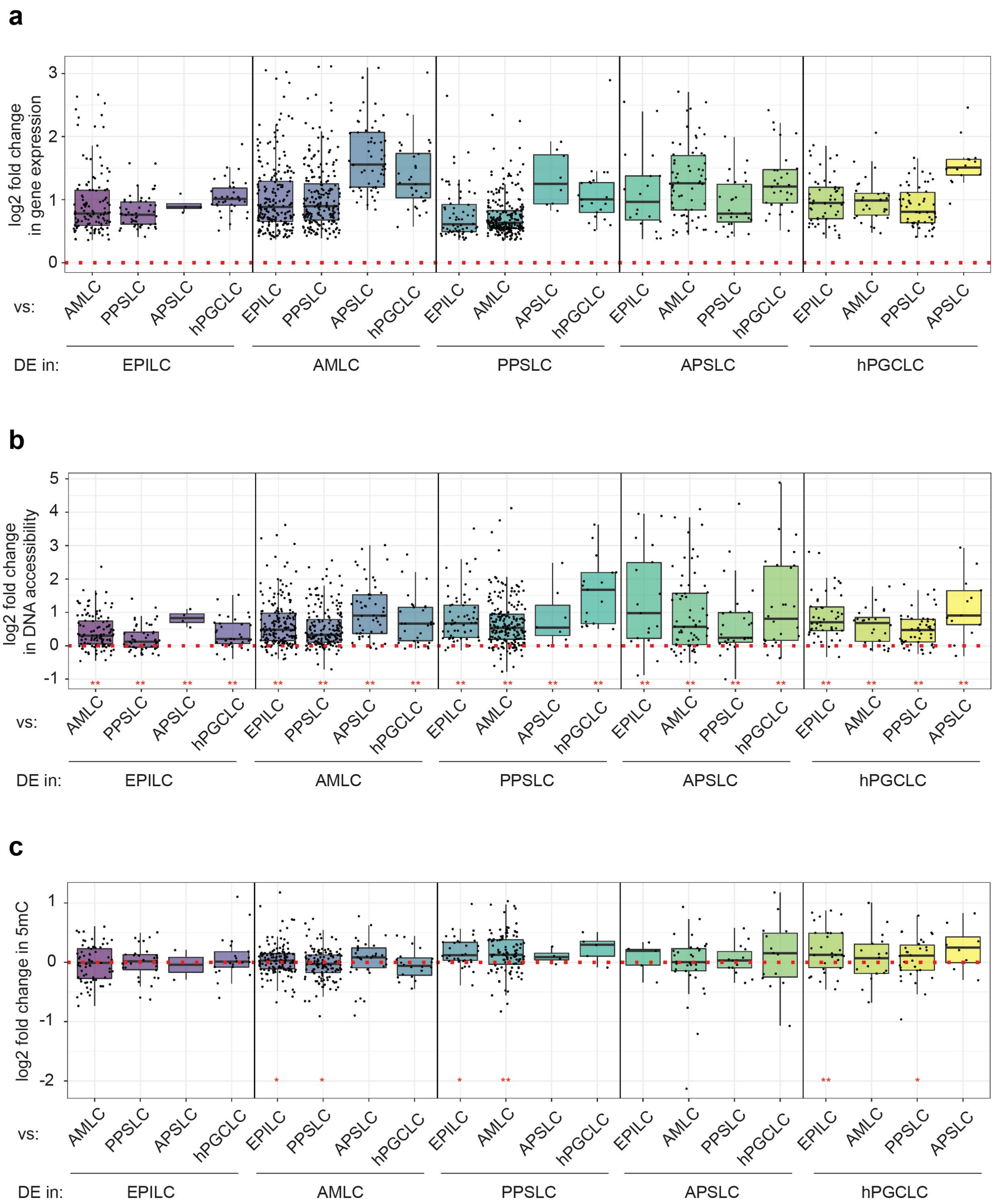


**Supplementary Figure 6 | The epigenetic landscape of cell types identified in the human gastruloids 48 hours post BMP4 addition.** (**a**) Log2 fold change in gene expression for DEGs in one cell type (DE in) compared to another cell type (vs). Note that the log2 fold change in mRNA is computed by taking the ratio of expression in a cell type where the gene is differentially expressed at a higher level compared to the other cell type. (**b**) Log2 fold change in DNA accessibility (promoter and gene body combined) for the genes depicted in (a). (**c**) Log2 fold change in gene body DNA methylation for the genes depicted in (a). Note that some of the genes shown in (a) are not depicted in (b) or (c) due to low detection across all cells. * and ** indicate a statistically significant change ($p < 0.05$ and $p < 0.01$, respectively, based on bootstrapped distributions from non-differentially expressed genes) in the log2 fold change of the epigenetic features for DEGs relative to non-differentially expressed genes.

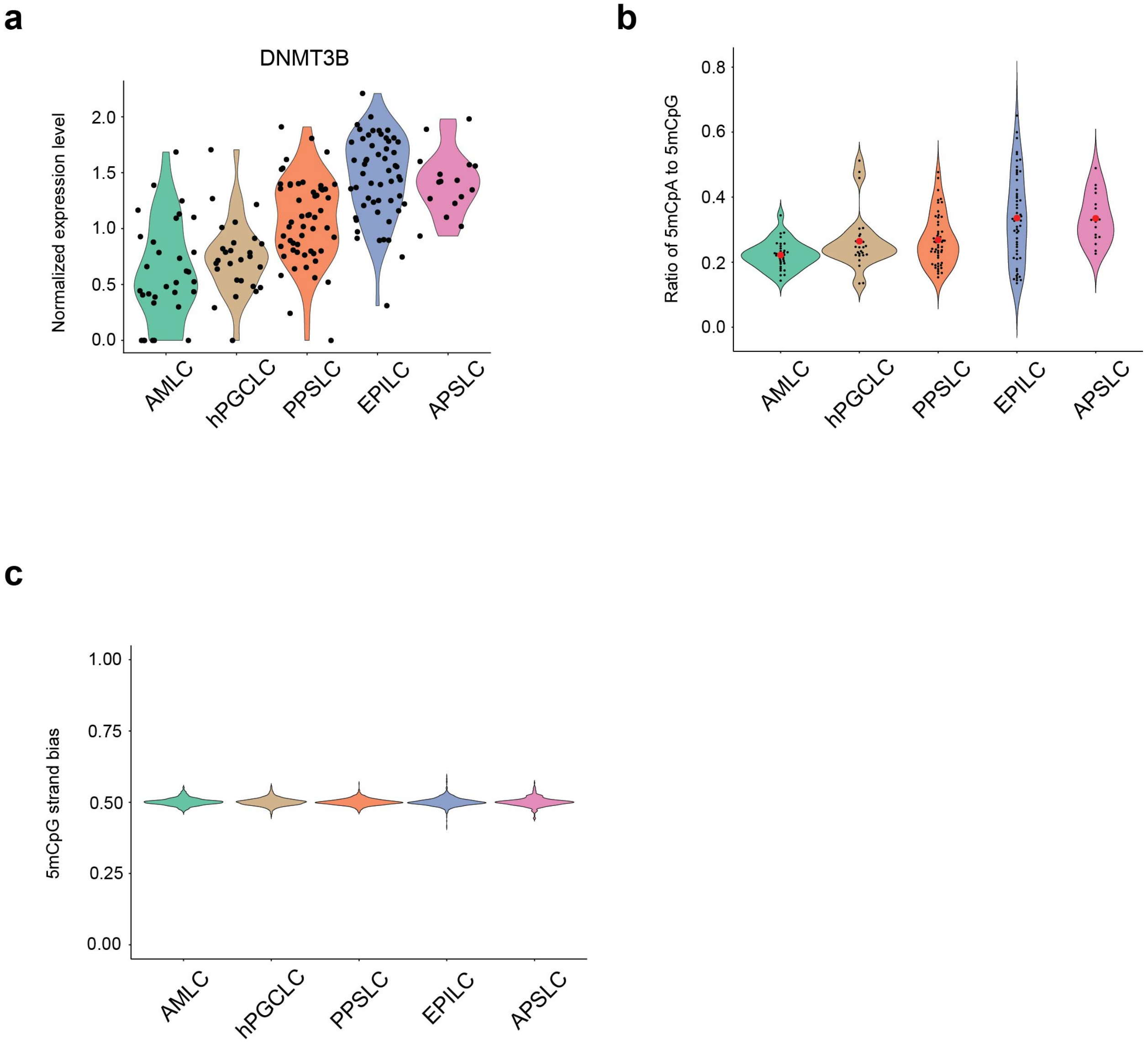


**Supplementary Figure 7 | DNA methylation dynamics in human gastruloids.** (**a**) Violin plot of gene expression levels for the *de novo* DNA methyltransferase DNMT3B for different cell types detected in the human gastruloid 48 hours post treatment with BMP4. Dots represent individual cells. (**b**) Violin plot of the ratio of 5mCpA to 5mCpG for different cell types. Black points represent individual cells, and the red point indicates the mean of the distribution. (**c**) Violin plot showing the distribution of 5mCpG strand bias in different cell types. 5mCpG strand bias is defined as the ratio of the number of 5mCpG sites detected on the plus strand of a chromosome over all 5mCpG sites detected on a chromosome. The panel shows that no 5mCpG strand bias is detected in any of the cell types.

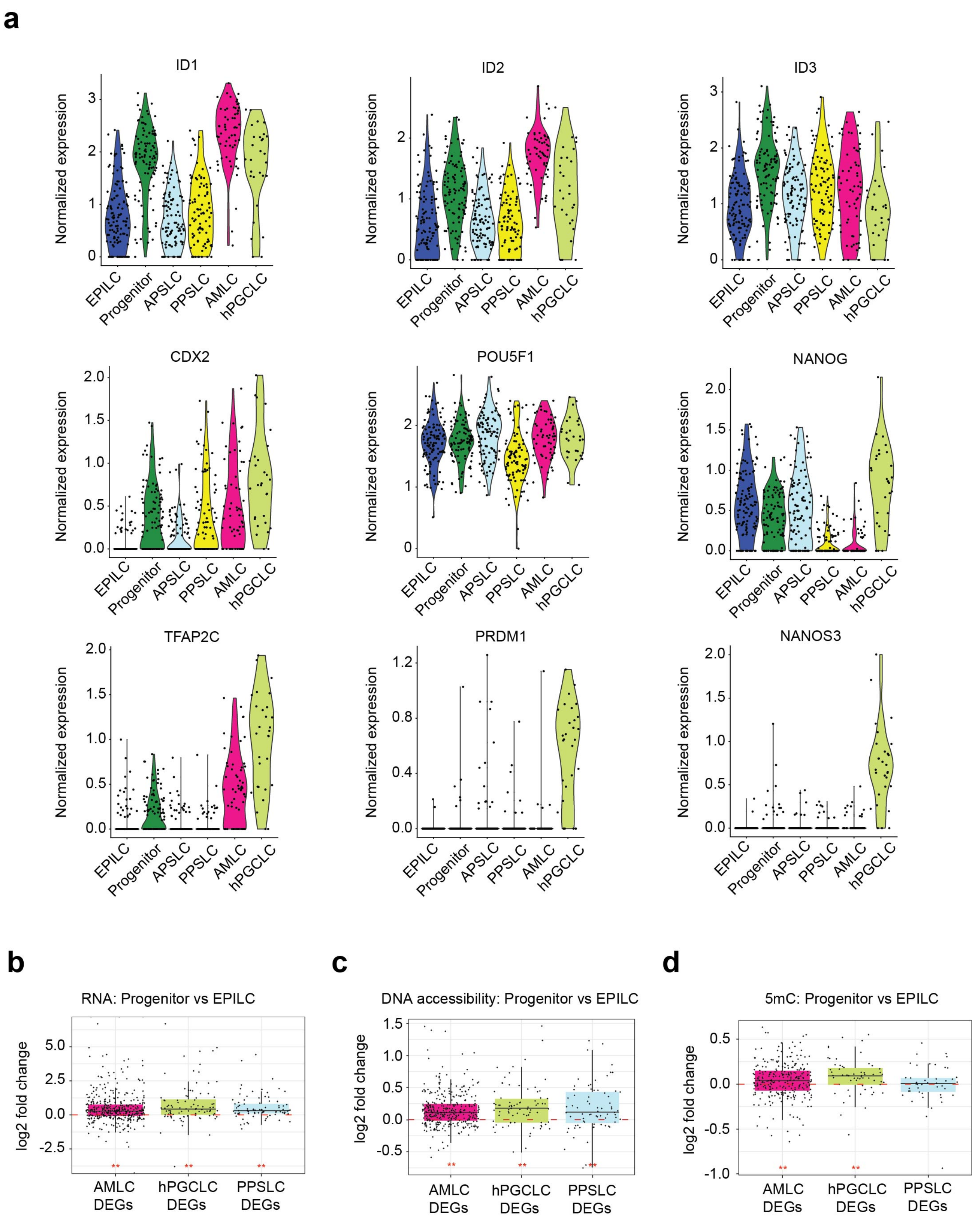


**Supplementary Figure 8 | hPGCLC and AMLCs bifurcate from a common progenitor population.** (**a**) Violin plots of gene expression levels of select genes for cell types identified in the human gastruloids at different timepoints after BMP4 addition. Genes shown here include targets of BMP4 signaling (ID1-3), an amnion related gene CDX2, genes related to pluripotency (POU5F1 and NANOG), and genes related to hPGCLC fate (TFAP2C, PRDM1, and NANOS3). (**b**) log2 fold change in gene expression for Progenitor cells compared to EPILCs for genes that are differentially expressed in the amnion (AMLC DEGs), hPGCLCs (hPGCLC DEGs) or PPSLCs (PPSLC DEGS) when compared to all other cell types. (**c**) log2 fold change in DNA accessibility (promoter and gene body combined) for the genes depicted in (b). (**d**) log2 fold change in gene body DNA methylation for the genes depicted in (b). ** indicates $p < 0.01$ (one-sample Wilcoxon signed rank test).

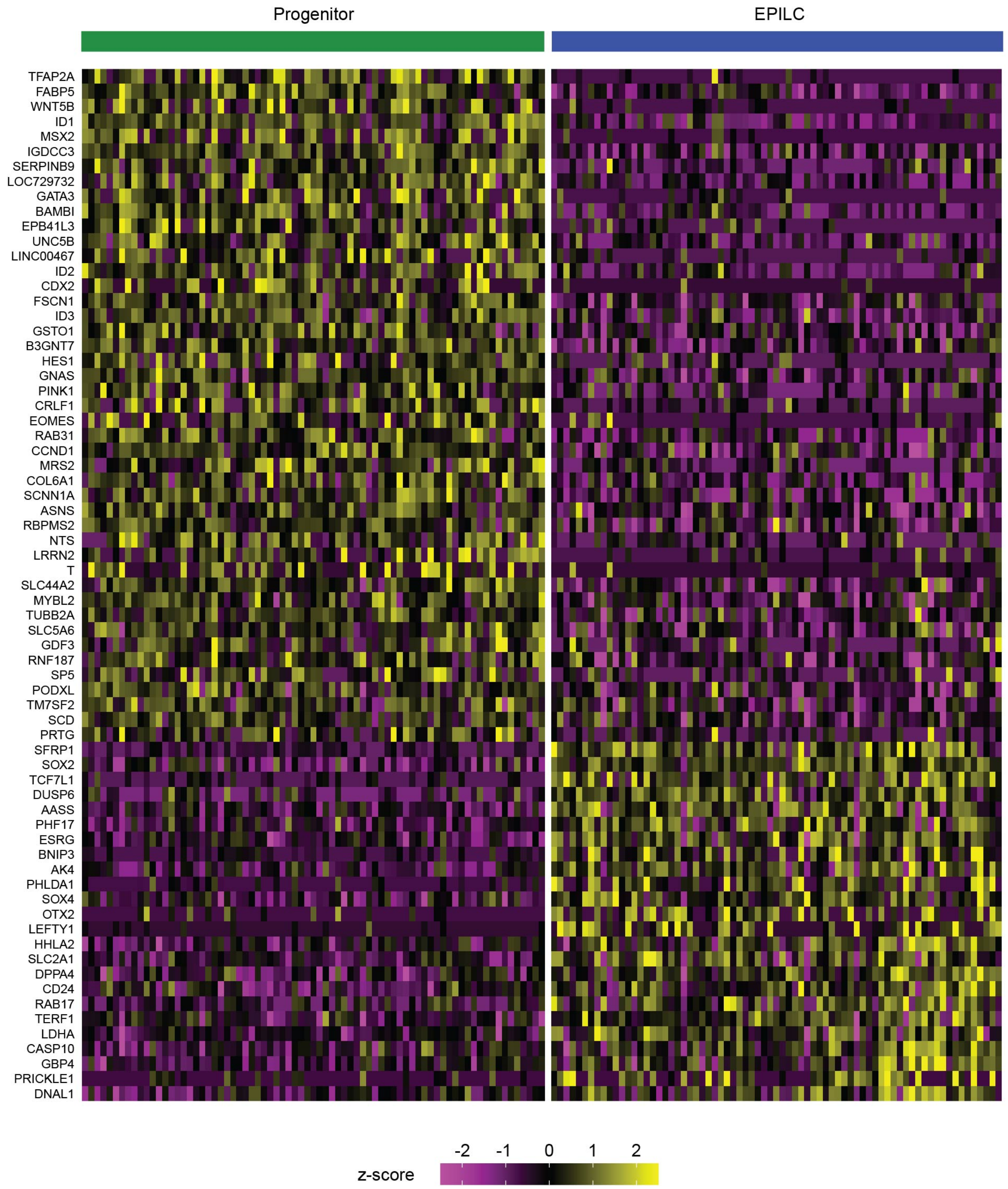


**Supplementary Figure 9 | DEGs between EPILCs and Progenitor cells.** Gene expression heatmap of z-scores for DEGs found between EPILC and Progenitor cells in human gastruloids 20 hours after treatment with BMP4.

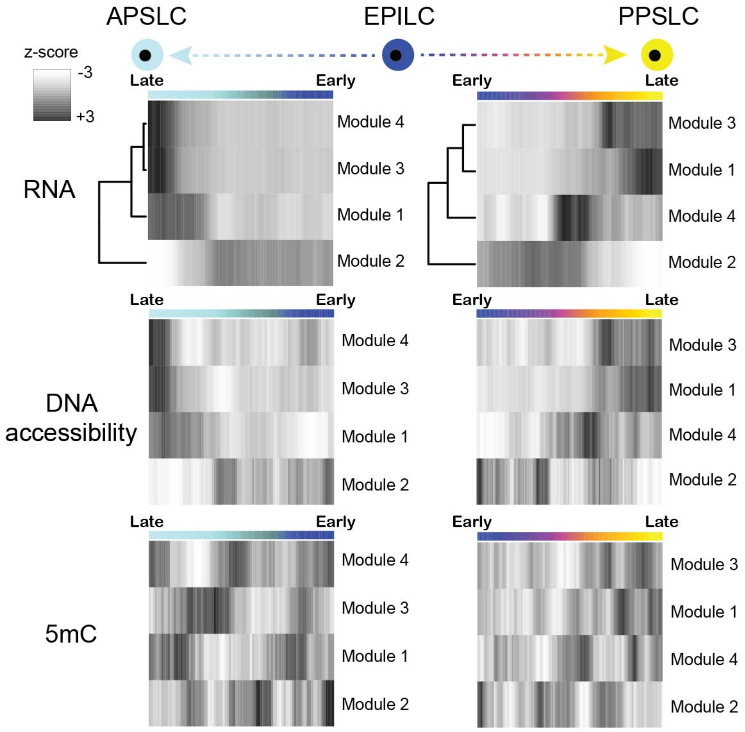


**Supplementary Figure 10 | APSLCs and PPSLCs arise from the EPILCs.** Heatmap of z-scores for DEG-derived gene modules along the APSLC and PPSLC pseudotime trajectory, together with the corresponding changes in DNA accessibility and DNA methylation. The color bar indicates position along the pseudotime, with EPILC in dark blue, APSLC in light blue, and PPSLC in yellow.
